# Trans-generational adaptation to maternal climate through hormone transport in plants

**DOI:** 10.1101/2024.10.04.616646

**Authors:** Xiaochao Chen, William Bezodis, Pablo González-Suárez, Vanda Knitlhoffer, Andrew Goldson, Ashleigh Lister, Iain Macaulay, Steven Penfield

## Abstract

Whether organisms can inherit parental adaptations to the environment is a major question in evolutionary biology. Plant development is highly plastic and dependent on the seasonal cues which are used to control growth and reproduction. Seed dormancy and germination are key traits which respond strongly to temperature during seed development and here we show that progeny adaptation to seasonal climate is inherited from the mother plant. Loss of maternal *LIKE HETEROCHROMATIN PROTEIN 1* (*LHP1*) causes an inability of progeny seeds to sense temperature and this is linked mechanistically to reduced ABA levels in seeds and activation of the primary nitrate response. At the single cell level, small changes in temperature activate nitrate signalling specifically in the mother, and ABA biosensor imaging reveals temperature-dependent fluxes of ABA into seeds necessary for dormancy induction. Thus, we reveal that progeny seeds inherit the climate adaptation of mother plants via active hormone transport during seed set.

## Introduction

Populations can adapt to environment through the action of natural selection on genetic and epigenetic variation. Epigenetic variation can also be induced by environmental changes and inherited into progeny (Carone et al., 2010; Klosin et al., 2017; Lin et al., 2024; Argaw-Denboba et al., 2024), but the extent to which progeny fitness is influenced by inheritance of parental adaptations to the environment remains unclear.

Plants are highly environmentally plastic, using seasonal environmental cues to align their development with time of year. Flowering time is tightly controlled to optimise the environmental conditions in which gamete production and seed set take place (Springthorpe and Penfield, 2015; Weitz et al., 2021). Cooler climatic conditions are associated with production of seeds with higher levels of dormancy, affecting the seasonal timing and frequency of germination (Von Abrams and Hand, 1956; Fenner, 1991; Postma and Agren, 2015; Iwasaki et al., 2022) and increasing the levels of the dormancy-inducing hormone abscisic acid (ABA) in mature seeds (Kendall et al., 2011).

Because effects of temperature on seeds are often accompanied by changes to maternal tissues such as fruit and seed coats (Li et al., 2017; Chandler et al,, 2024; Chen et al., 2014; Fedi et al., 2017) it has long been speculated that effects of temperature are transmitted trans-generationally from the mother plant (Roach and Wulff, 1987). Maternally inherited epialleles may be important for effects of environmental variation on seeds (Iwasaki et al., 2019) but cannot account for the strong effects of the post-fertlisation environment on seed behaviour.

LIKE HETEROCHROMATIN PROTEIN 1 (LHP1), the plant orthologue of mammalian HETEROCHROMATIN 1, interacts with POLYCOMB REPRESSIVE COMPLEX 2 (PRC2) and is required for the formation of facultative heterochromatin in plants (Derkecheva et al., 2013). Here we show that disruption of LHP1 in mother plants prevents seed dormancy induction by low temperatures. However, maternal eipalleles do not need to be inherited into progeny, but act through constitutive de-repression of the Primary Nitrate Response (PNR; Krapp et al., 2014) and changes to ABA levels in the mother plant. Remarkably, we show that warm temperatures also activate the PNR in mother plants and control dormancy via trans-generational flux of ABA into progeny seeds. Thus, ABA is a trans-generational signal through which plant populations inherit maternal adaptation to seasonal climate.

## Results

Previously, we have shown that lowering the temperature from 22°C to 16°C during Col-0 seed set is sufficient to induce seed dormancy. We found that at 22°C developing seeds already showed detectable low dormancy 12 days after pollination and develop high germination during later stages of seed maturation (Figure 1A). At 16°C we could not detect germination at any time during seed development, suggesting that temperature affects dormancy throughout seed development. Previously, we showed that the PRC2-accessory protein VERNALISATION INDEPENDENT 3-LIKE 3 (VEL3) acts in the central cell to establish imprinted genes in the endosperm necessary for seed dormancy induction by low temperatures (Chen et al., 2023). Because LHP1 acts alongside PRC2 in transcriptional repression we tested whether *Arabidopsis* LHP1 has a role in low-temperature induction of seed dormancy. We found that *lhp1-6* mutants showed a non-dormant germination pattern during seed development at 16°C that was qualitatively similar to that observed in WT at 22°C, although seed development remained delayed by low temperature (Figure 1A). This showed that LHP1 is required for seed dormancy responses to temperature, but not for the temperature-induced delay to seed development. This phenotype was confirmed in further *lhp1* mutant alleles (Figure S1A) and could be complemented by expression of LHP1-GFP using the endogenous promoter but not when using the endosperm-specific EPR promoter (Figure S1B).

**Figure 1.**
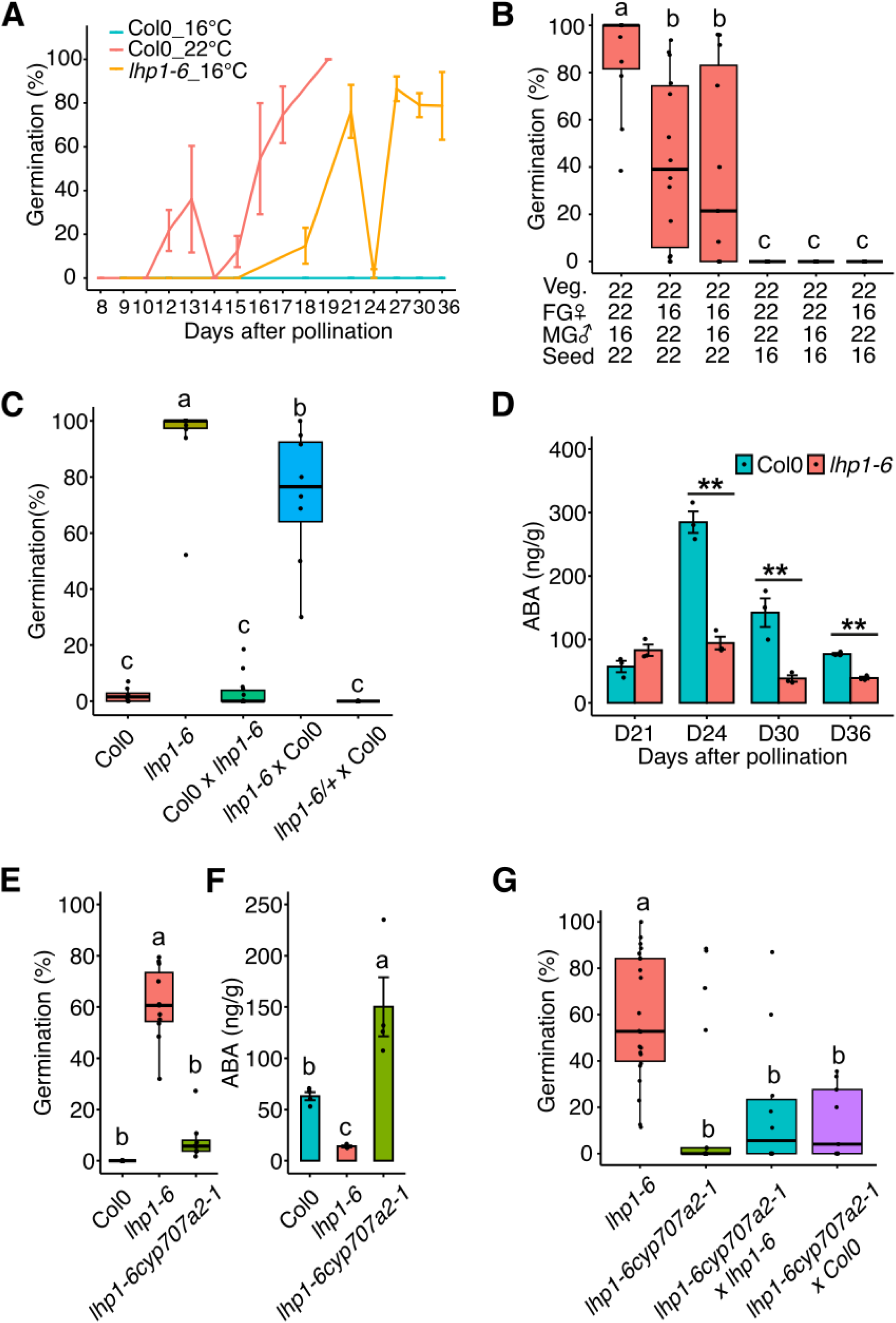
Temperature regulation of progeny seed dormancy is a maternal phenomenon that requires LHP1-regulation of ABA levels. **A**. Seed dormancy was tested by germination of seeds dissected from developing siliques throughout seed development, showing that *lhp1-6* plants develop low dormant seeds at cool temperatures, and that in WT temperature acts early in seed development to affect dormancy. Images showing seed morphology at corresponding stages are shown in Figure S1F. Data shows mean +/- SE, n=3-5. **B.** Temperature experience of the mother plant affects progeny dormancy. Germination of freshly harvested seed from plants given 16°C or 22°C before or after flowering, and seeds were generated from crosses using mother plant (MG) or father plants (FG) growing at different temperatures before flowering as indicated (Veg. = Vegetative Growth, MG = Male Gametophyte, FG = Female Gametophyte) . Significant differences are shown using Tukey post hoc test (P < 0.05; n = 10-16). **C**. LHP1 is required in the maternal sporophyte to affect seed dormancy. Dormancy was tested by germination of progeny seeds generated at 16°C using reciprocal crosses between WT and *lhp1-6*. Data is shown is from 8-14 biological replicates per genotype. Maternal genotype is given first for F_1_ seeds from crosses. **D**. LHP1 is required for ABA accumulation in seeds at 16°C. ABA was measured using LC-MS at the indicated timepoints (D – Day). At D36 WT seeds reach maturity. Significant differences are shown (P < 0.01 n= 3). **E**. Disruption of ABA catabolism restores normal seed dormancy in *lhp1-6*. Comparison of germination frequency of freshly harvested WT and *lhp1-6* and *lhp1-6 cyp707a2-1* double mutant seeds set at 16°C. Significant differences are shown (P < 0.05 n= 8-12). **F**. The low ABA level of *lhp1-6* seeds is increased in *lhp1-6 cyp707a2-1* seeds set at 16°C, as measured in mature dry seeds by LC-MS. Significant differences are shown (P < 0.01 n= 4). **G**. For *lhp1-6 cyp707a2-1* the maternal genotype of *CYP707A2* is important for the rescue of ABA levels in the double mutant. Dormancy was measured by germination of freshly harvested seeds set at 16°C, maternal genotype is given first in seeds derived from crosses. Significant differences are shown (P < 0.01 n= 9-22).

To determine whether parent plants play a role in the effect of temperature on seed dormancy, we set seed from reciprocal crosses between mothers and fathers grown at either at 22°C or 16°C. This revealed that the pre-fertilisation temperature effect on progeny seed dormancy is maternally inherited, because changing the paternal growth temperature had no impact on progeny dormancy (Figure 1B). To test whether LHP1 is required for temperature-regulation of progeny seed dormancy by the mother, we analysed dormancy of progeny of reciprocal crosses between *lhp1-6* and WT at 16°C (Figure 1C). This confirmed that *lhp1-6* abolishes seed dormancy responses to temperature: furthermore, we found that loss of LHP1 in the maternal sporophytic tissue is necessary and sufficient to observe a strong phenotype. Therefore we concluded that temperature responsive seed dormancy, caused by temperature variation during seed development, is also explained primarily by a maternal effect that is largely abolished in progeny of *lhp1* mutant mother plants.

Induction of seed dormancy by low temperatures requires an intact seed coat which is derived from the maternal ovule integuments (MacGregor et al., 2015; Fedi et al., 2019) and is associated with an increase in abscisic acid (ABA) levels in the seed (Kendall et al., 2011; Iwasaki et al 2019). Our analysis of *lhp1* seed coats suggested that these resembled WT in colour and permeability (Figure S1C). However, we found that *lhp1* mutant seeds set at 16°C showed consistently lower ABA levels, which were maintained until the mature dry state (Figure 1D). Furthermore, crossing *lhp1-6* to the ABA-hyperaccumulating *cyp707a2-1* mutant raised ABA levels in mature seeds and restored seed dormancy (Figure 1E, F). F_1_ seeds from *lhp1-6 cyp707a2-1* double mutant mother plants crossed to WT also showed normal dormancy induction by low temperatures (Figure 1G). This reduced ABA level seed dormancy in *lhp1-6* compared to Col-0 is also observable at 22°C (Figure S1D,E) and supports an additive effect of *lhp1* loss and temperature. We therefore concluded that loss of *LHP1* in mother plants abolishes progeny seed dormancy responses to temperature via regulation of ABA levels: our data (Figure 1G) suggested that maternal ABA plays a role in this process.

To further examine the importance of maternal tissues in seed dormancy we used a fruit culture system (Creff et al., 2018; see methods) setting viable seeds in liquid culture (Figure S2A), for seed dormancy testing. We noticed that during fruit culture at 16°C, *lhp1-6* mutants would undergo pre-harvest sprouting, unlike WT seeds which did not germinate in culture while still attached to the replum (Figure S2B,C). However, WT seeds that detach from the fruit germinated in the culture medium, likely because 16°C induces a cold-stratification response (Springthorpe and Penfield, 2015). Thus developing seeds themselves are not able to induce dormancy, but for the duration of the fruit culture presence of fruit tissue is sufficient. So we focused on analysing LHP1-dependent processes in fruits.

To understand the mechanism by which loss of maternal LHP1 affects seed dormancy we performed RNAseq in fruit tissues at 14 DAP at 16°C, removing the seeds. GO-term analysis of genes up-regulated in *lhp1-6* fruits revealed that the most over-represented category was ‘response to nitrate’, suggesting that *lhp1* mutants have constitutively active nitrate signalling. Further analysis showed that this includes most of the major genes forming the so-called Primary Nitrate Response (PNR; Shanks et al., 2020; Krapp et al., 2014; Figure 2A, B). Among the most strongly *lhp1* down-regulated genes was *NCED3*, the rate limiting step in ABA synthesis, alongside transcription factors with known roles in *NCED3* regulation during lateral bud dormancy such as *HB40* and *NAC29/NAP* (Figure 2C), a response that is also repressed by nitrate supply (González-Grandío et al., 2017; de Jong et al., 2014). This raised the question of whether the elevated PNR in *lhp1-6* mutants was accompanied by changes to tissue nitrate levels and we found that during flowering and seed set *lhp1* mutants accumulate high nitrate levels throughout the plant including the rosette leaves, fruits and seeds (Figure 2D). Interrogating a previous analysis of LHP1 genome-wide binding sites (Veluchamy et al., 2016) revealed few nitrate-responsive genes with strong LHP1 binding signals: however, two nitrate transporters, *NRT1.5* and *NRT2.1* are clearly direct targets of LHP1. Loss of either transporter gene reduced seed nitrate levels and led to lower germination rates of progeny seeds (Figure 2E-G).

**Figure 2.**
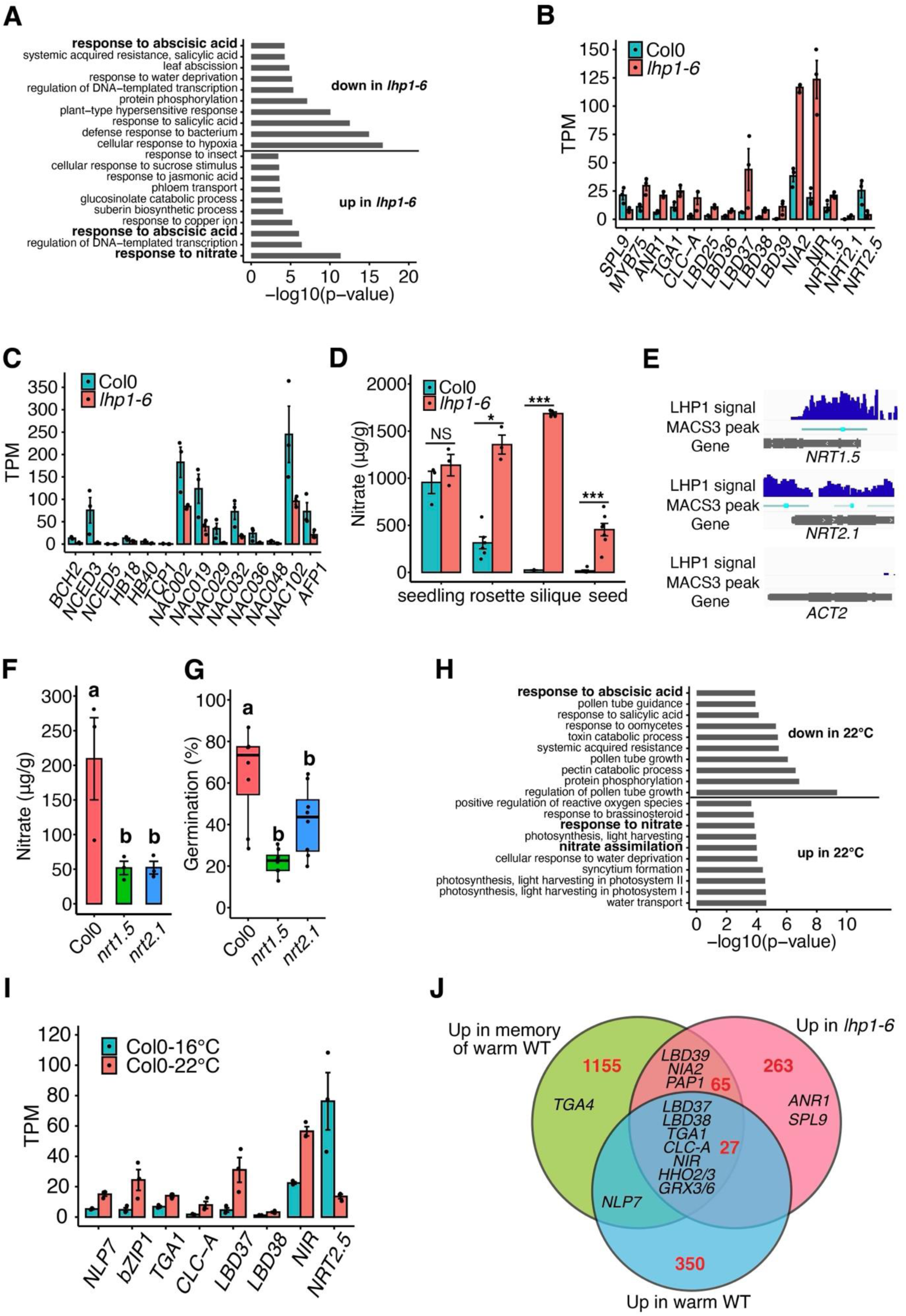
The effect of maternal *lhp1* on dormancy is mediated by nitrate accumulation and signalling. **A.** GO term analysis of differentially expressed genes (DEGs) in fruits that are upregulated or downregulated in *lhp1* vs WT fruits. Full list of DEGs and GO terms in Table S1 and S2. **B, C.** RNAseq data showing effects of LHP1 loss on the primary nitrate response (B) and ABA synthesis (C). **D.** Nitrate hyperaccumulation in *lhp1* plant tissues in seedlings or various tissues during seed set. Data shows mean +/- SE, n = 3-7. **E** ChIP-seq data reveals that nitrate transporters are direct targets of LHP1. Data reanlysed from Velucharmy et al. (2016), compared to ACTIN2 negative control. **F.** Nitrate levels in WT, *nrt1.5* and *nrt 2.1* freshly harvested dry seeds set at 22°C. Data show mean +/- SE, n = 3. Significant differences shown by Tukey post hoc test (P<0.05). **G**. NRT1.5 and *NRT2.1* are necessary for low dormancy at 22°C. Germination frequency of freshly harvested seeds, significant differences shown by Tukey post hoc test (P<0.05). **H.** GO-Term analysis of DEGs in fruits that are upregulated or downregulated by temperature shows that temperature changes affect expression of PNR and ABA responses. **I.** Effect of temperature on expression of individual PNR-associated DEGs in fruits. **J.** Upregulation of well-characterised nitrate responsive genes shared between *lhp1* vs WT at 16°C, WT 22°C vs 16°C and WT (Ler) at 22°C vs WT (Ler) at 22°C but grown until flowering at 16°C (memory of cold), latter data from Chen et al. (2014).

Because nitrate availability affects progeny seed dormancy through regulation of ABA metabolism (Matakiadis et al., 2009), de-regulation of the primary nitrate response in *lhp1* mutants could over-ride the effect of low temperature on progeny dormancy by repressing ABA production. However, a second possibility was that temperature itself could act via an effect on the PNR. To test this, we also compared fruit transcriptomes at the bent cotyledon stage developing since fertilisation at 22°C or 16°C (Figure 2H-J). Remarkably, in fruits the effect of the temperature increase resembled loss of *LHP1*, with warm temperatures inducing genes in early nitrate assimilation and well-characterized transcription factors such as *ANR1*, *TGA1*, *LBD37-39* and *NLP7*. We also cross-referenced this dataset to a previously obtained RNAseq dataset comparing gene expression in fruits from plants which experienced similar high and low temperature treatments before rather than during flowering and seed set (Chen et al., 2014; hereafter referred to as memory of warm). Interestingly of the 27 genes commonly upregulated in fruits in *lhp1* mutants at 16°C, WT at 22°C and WT memory of 22°C versus WT at 16°C many were part of the PNR including *TGA1*, *NIR*, *CLC-A*, *LBD37/38* and *HHO2/3*, while *NLP7*, *LBD39*, *PAP1*, and *NIA2* were up-regulated in two of the three experiments (Figure 2J). This suggested the hypothesis that temperature affects seed dormancy via activation of the PNR, either in the mother plant or developing seeds.

Fruits and seeds are morphologically highly complex, containing mixtures of maternal tissues and zygotic embryo and endosperm tissues with differing genotypes. To further understand the tissue-specific distribution of ABA, nitrate and temperature responses in developing seeds we conducted single nucleus RNAseq using the 10X Chromium platform. We isolated nuclei from whole siliques at the bent cotyledon stage of seed development, from Col-0 at 16°C and 22°C and *lhp1-6* mutant siliques at 16°C. To ensure developmental stages were closely matched, the 22°C treatment was grown at 16°C until one day before sampling and subjected to the warmer temperature treatment for 24 hours. A previous single cell analysis of whole siliques divided the resulting nuclei into 26 major clusters (Lee et al., 2023). This is similar to the 22 clusters we obtained using clustering at Seurat clustering resolution 1 (Figure S3A). As we wished to locate key responses in readily identifiable cell types, we chose a clustering resolution to give us 13 clusters with a clear maternal and zygotic separation in which major tissue types such as seed coat cell types, funiculus and fruit vascular tissues that arise primarily from the placenta could be identified using well characterised marker genes (Figure 3A, B; Table S3). These were then independently confirmed using Differentially Expressed Genes (DEGs) comparing transcriptomes from whole fruits and seeds (Table S4) and a pre-existing laser capture microdissection study (Belmonte et al. 2013; Figure S3B). Further reducing the clustering resolution clearly divided out samples into one maternal super-cluster containing all fruit and seed coat tissues, one for endosperm and one for embryo indicating that genotype is the most important driver of gene expression differences between fruit and seed tissues (Figure S3A).

**Figure 3.**
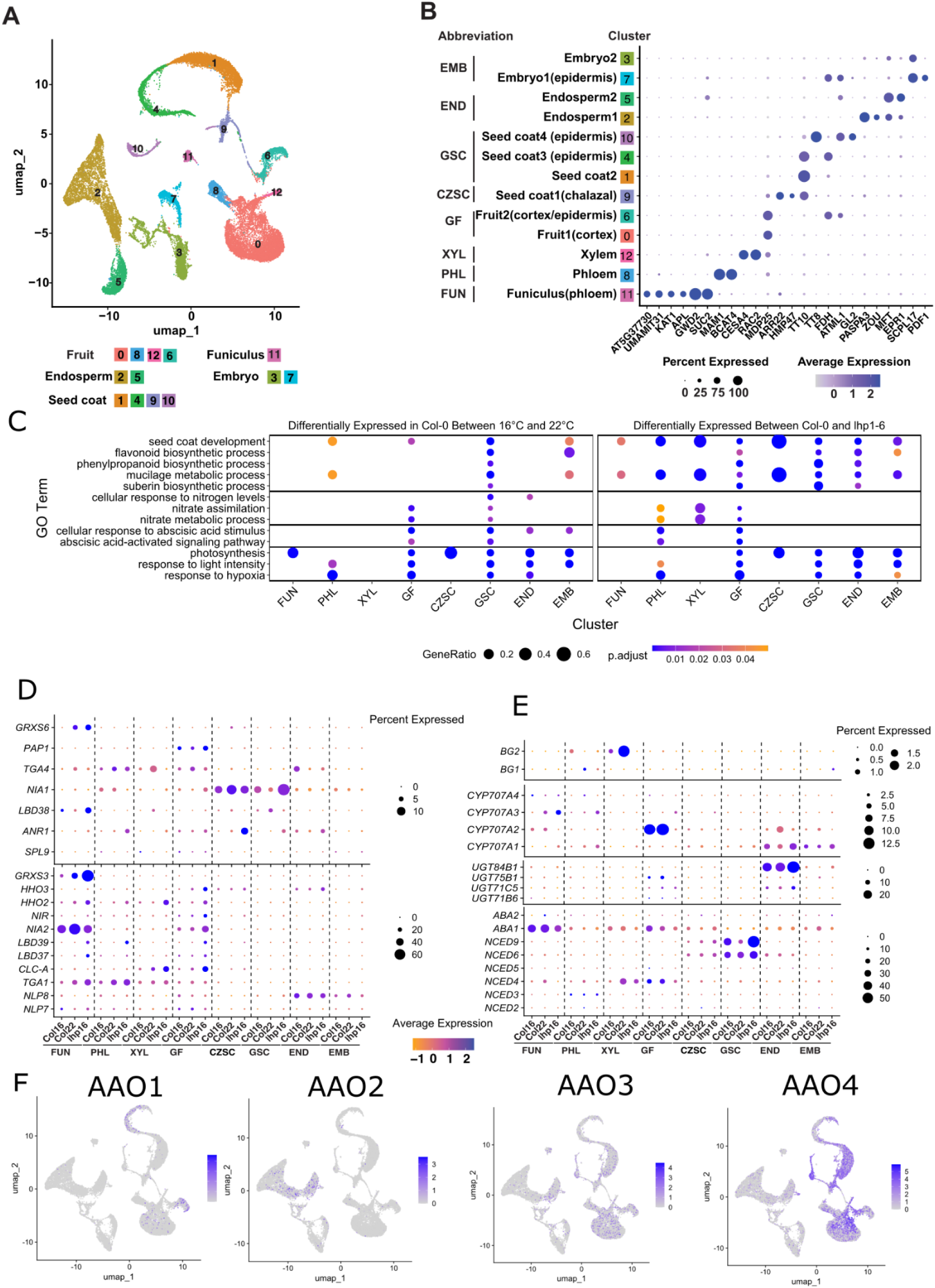
snRNA-seq of developing fruits including seeds reveals tissue-specific patterns of ABA metabolism and nitrate responses. **A.** Integrated UMAP plot of nuclei, coloured by cluster with tissue types noted, based on the annotation in B. **B.** Expression of well-characterised marker genes shows the identity of each cluster with broader tissue types indicated (FUN = Funiculus, PHL = Phloem, XYL = Xylem, GF = General Fruit, CZSC = Chalazal Zone of Seed Coat, GSC = General Seed Coat, END = Endosperm, EMB = Embryo). Size of circles indicates the percentage of cells in that cluster expressing each marker gene and colour indicates expression levels. **C.** Analysis of selected GO Terms across clusters based on cluster-specific DEGs comparing the effect of temperature on WT fruits and comparing WT and *lhp1-6*. Sizes of circles indicates the Gene Ratio and shading indicates adjusted p-values of GO enrichment. Complete lists of GO terms and DEGs for each cluster comparison is provided in supplementary tables S5 and S6. **D.** Dotplot showing expression of key genes involved in nitrate perception or response by tissue type, temperature and genotype . **E.** Similar Dotplot with key genes involved in ABA biosynthesis, degradation, conjugation to ABA-GE and release of ABA from ABA-GE. **F** UMAP plots of showing expression of *AAO1-4*, the final step in bioactive ABA synthesis showing that expression is primarily in maternal tissues (clusters labelled in panels A and B).

We identified DEGs in each major cluster (Table S5) and used GO-terms to identify biological processes associated with different tissue types (Table S6). Firstly, we found that gene expression relating to nitrate assimilation and metabolism were affected by temperature only in the maternal tissues of the fruit and seed coat (Figure 3C), whereas changes in the response to nitrate were seen in seed coat and endosperm. In *lhp1-6* mutants, effects on nitrate metabolism were confined to fruit tissues. A similar pattern was seen for ABA responses. Temperature and LHP1 had similar effects on photosynthetic and light intensity responses which were highly activated by either warm temperatures or LHP1 across a range of tissues. More detailed analysis of the PNR revealed cell type-specific responses of that were particularly strong in *lhp1-6* mutants, including *LBD37* and *LBD39* in the funiculus and fruit phloem, *CLC-A* in fruit mesophyll and fruit vascular cells, *TGA1* in all fruit tissue clusters and *ANR1* in fruit phloem and the chalazal seed coat. The exception was *NLP8* expression which was detected in zygotic tissues, consistent with its role in nitrate responses of imbibed seeds after shedding (Yan et al., 2016) but was not affected transcriptionally by our treatments. In general PNR responsiveness was weak or absent from embryo and endosperm clusters, demonstrating that at the molecular level the effect of LHP1 and temperature on nitrate responses is characteristic of maternal tissues, and despite the fact that *lhp1-6* seeds accumulate elevated nitrate (Figure 2D).

Whole tissue RNAseq indicated that the biggest effect of *lhp1* on ABA metabolism was on *NCED3* expression (Figure 2). *NCED3* was mainly expressed in the phloem of the fruit suggesting that the vascular tissue is a primary site of ABA synthesis in low nitrate conditions (Figure 3E). The early steps in the ABA synthesis pathway, *ABA1* and *ABA2*, were expressed constitutively at similar levels in all tissue types, while *NCEDs*, *AAOs* and *CYP707A*-encoding genes were expressed in a non-overlapping tissue specific manner. *NCED6* and *NCED9* were detected in the seed coat and chalazal seed coat, although these were most highly expressed in the non-dormant *lhp1-6* mutant (Figure 3E). The last step in ABA synthesis is catalysed by ABSCISIC ALDEHYDE OXIDASE (AAO). We mined data for the expression of *AAO1-4* gene expression which were found almost exclusively in maternal tissues of the fruit and seed coat, confirming the dominance of maternal ABA synthesis at this developmental stage relative to the zygote (Figure 3F). However, *NCED*, *AAO or CYP707A* expression were not clearly affected by temperature in fruits and seeds which left the source of temperature-regulated differences in ABA unclear.

Our data suggested that effects of temperature are associated with impacts on nitrate responses, but previous work suggests there is no clear relationship between temperature and nitrate accumulation in seeds (Huang et al., 2018), a finding we could confirm (Figure 4A). However, we found that feeding excess nitrate to mother plants during seed set at 16°C could strongly counteract the effect of low temperature on progeny dormancy (Figure 4B, C). Two more experiments confirmed that nitrate given to mother plants was able to substitute for the temperature signal: firstly, we set seeds using cut inflorescences in culture medium (see methods) at either 22°C or 16°C. High nitrate feeding resulted in production of non-dormant seeds at either temperature, but under lower nitrate feeding, lower temperatures induced dormancy (Figure 4D). A third experiment exploited seed set in fruit culture: in this system we confirmed that high nitrate feeding resulted in non-dormant seeds regardless of fruit culture temperature, whereas elimination of nitrate resulted in large fractions of dormant seeds in both the warm and the cold (Figure 4E). Under intermediate nitrate feeding the normal effect of maternal temperature on dormancy could be observed. Because we could not separate the temperature signals from the effect of nitrate itself, we measured whether changing temperature affects nitrate concentrations of plant tissues during reproduction. In rosette leaves, nitrate levels fell during reproductive development, but lower temperatures consistently reduced nitrate levels relative to warm conditions in flowering plants. A broadly similar trend was observed in inflorescence tissues (Figure 4F, G; Figure S6). Therefore, we concluded that affects of temperature on nitrate responses are sufficient to impact progeny dormancy.

**Figure 4.**
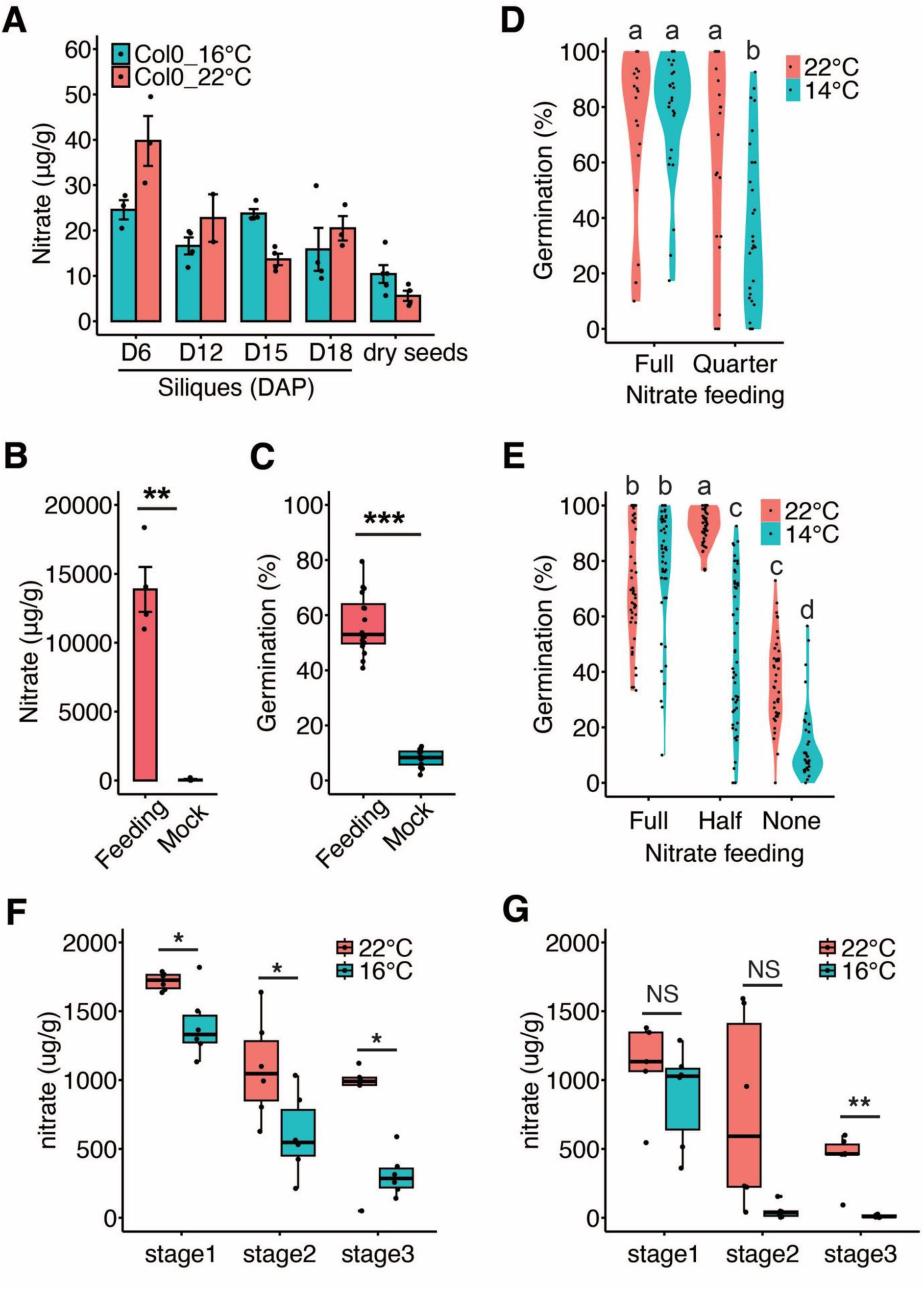
In WT Col-0, Nitrate levels are correlated with temperature and maternal supply of nitrate leads to reduced dormancy. **A.** Nitrate levels in developing fruits (including seeds) and mature seeds are not mainly unaffected by temperature. Charts show mean +/- SE n= 3-4 . **B.** Excess maternal feeding of nitrate leads to nitrate hyperaccumulation in dry seeds. Charts show mean +/- SE n= 4. **C.** Germination of freshly harvested seeds set at 16°C with or without high nitrate supply to the mother plant. Charts show mean +/- SE n= 16 . **D.** Fruit culture with reduced maternal supply of nitrate induces seed dormancy only at low temperatures: gemination frequency of freshly harvested mature seeds from fruit culture at the indicated temperature and nitrate feeding levels. Significant differences are shown by Tukey post hoc test at P < 0. 05, n= 22-30). **E.** Germination of freshly harvested seeds set in inflorescence cultures in the indicated temperature and nitrate regimes. Significant differences are shown by Tukey post hoc test at P < 0.05, n = 37-50. **F.** and **G.** Nitrate levels in flowering plants grown at low or high temperature at matched developmental stages in rosette leaves (**F**) and inflorescence stems (**G**). Stages are described in Figure S4.

Temperature affects nitrate levels in tissues of the mother plant but not in fruits and seeds, suggesting that nitrate itself is not the trans-generational signal. This conclusion is further supported by restriction of nitrate responses to the fruit and seed coat tissues in single cell RNAseq data (Figure 3). In the case of *lhp1-6* mutants we show that this ABA is also maternal origin, (Figure 1G) so we tested whether maternal ABA is necessary and sufficient for low temperatures to induce dormancy. Importantly, application of ABA to imbibed seeds prevents seedling establishment but cannot induce dormancy (Lopez-Molina et al., 2002; Penfield et al., 2006). However, feeding increasing ABA concentrations to fruits cultured at 22°C or 14°C resulted in seeds with progressively higher dormancy levels at maturity (Figure 5A). Thus, ABA applied to mother plants is sufficient to induce dormancy in progeny seeds. In these experiments, ABA accumulated in seed tissues at higher levels than in the culture media (Figure 5B), suggesting that fruits concentrate ABA in seeds via active transport. To rule out that supplied ABA was activating *de novo* ABA synthesis in seeds, we also fed deuterated ABA (d6-ABA) to detached inflorescences: both ABA and deuterated d6-ABA could be detected in seeds demonstrating ABA transport throughout the inflorescence to the seeds (Figure 5C). Next, we tested whether maternal ABA was necessary for progeny seed dormancy at 16°C using reciprocal crosses between WT and the *aba2-1* mutant, and between heterozygous mothers and WT (Figure 5D). These revealed that maternal ABA accounted for the majority of dormancy induction by low temperatures and this effect is of maternal sporophytic origin. Thus, maternal ABA is necessary and sufficient for seed dormancy responses to temperature.

**Figure 5.**
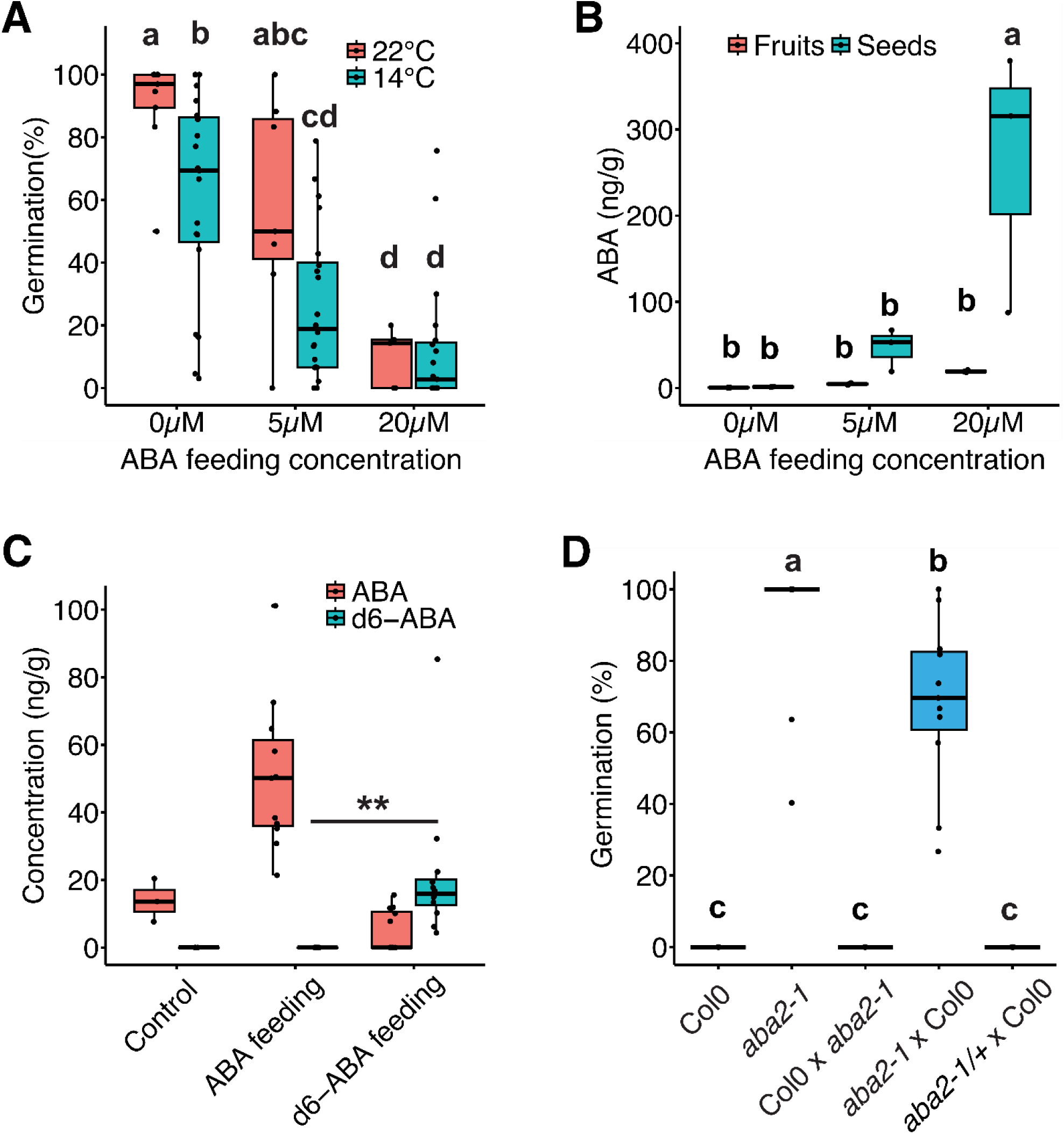
ABA supplied maternally or exogenously during development is necessary and sufficient for low temperature to induce seed dormancy. **A.** Germination of freshly harvested seeds with or without added of ABA to fruit cultures at the indicated temperatures. Chart shows mean +/- SE n= 5-20 and significant differences shown by Tukey post hoc test at P < 0. 05. **B.** LC-MS measurements of ABA levels in fruit and seed produced after fruit culture as in (A) showing accumulation of high levels of ABA in seed tissues (n = 3). **C.** LC-MS detects exogenously supplied d6 ABA to the base of inflorescences in seeds. n = 3-12, significant differences shown by t.test (P < 0.05). **D.** Germination of freshly harvested WT and *aba2-1* seed set at 16°C and resulting reciprocal crosses. Significant differences shown by Tukey post hoc test (P < 0.05, n = 11-12).

Analysis of cell type specific *AAO* expression revealed that the final step of ABA synthesis takes place in fruit cells and fruit phloem (Figure 3F). To test whether changing temperature increased ABA levels in fruits and seeds we measured ABA in whole fruit tissues and developing seeds at bent cotyledon-mature green stage by LC-MS. Lowering the temperature from 22°C to 16°C for 24 hours clearly increased ABA levels in seeds but fruits showed no substantial changes (Figure 6A). To understand in more detail how temperature affects ABA dynamics in individual tissues we exploited the ABACUS2 ABA biosensor (Rowe et al., 2023), imaging developing fruits at either 14°C or 22°C (Figure 6B-E). Imaging seeds within fruits at 14°C we observed an apparent gradient in ABA levels in developing seeds with highest ABA at the chalazal pole and funiculus, suggestive of ABA transport into seeds (Figure 6B).

**Figure 6.**
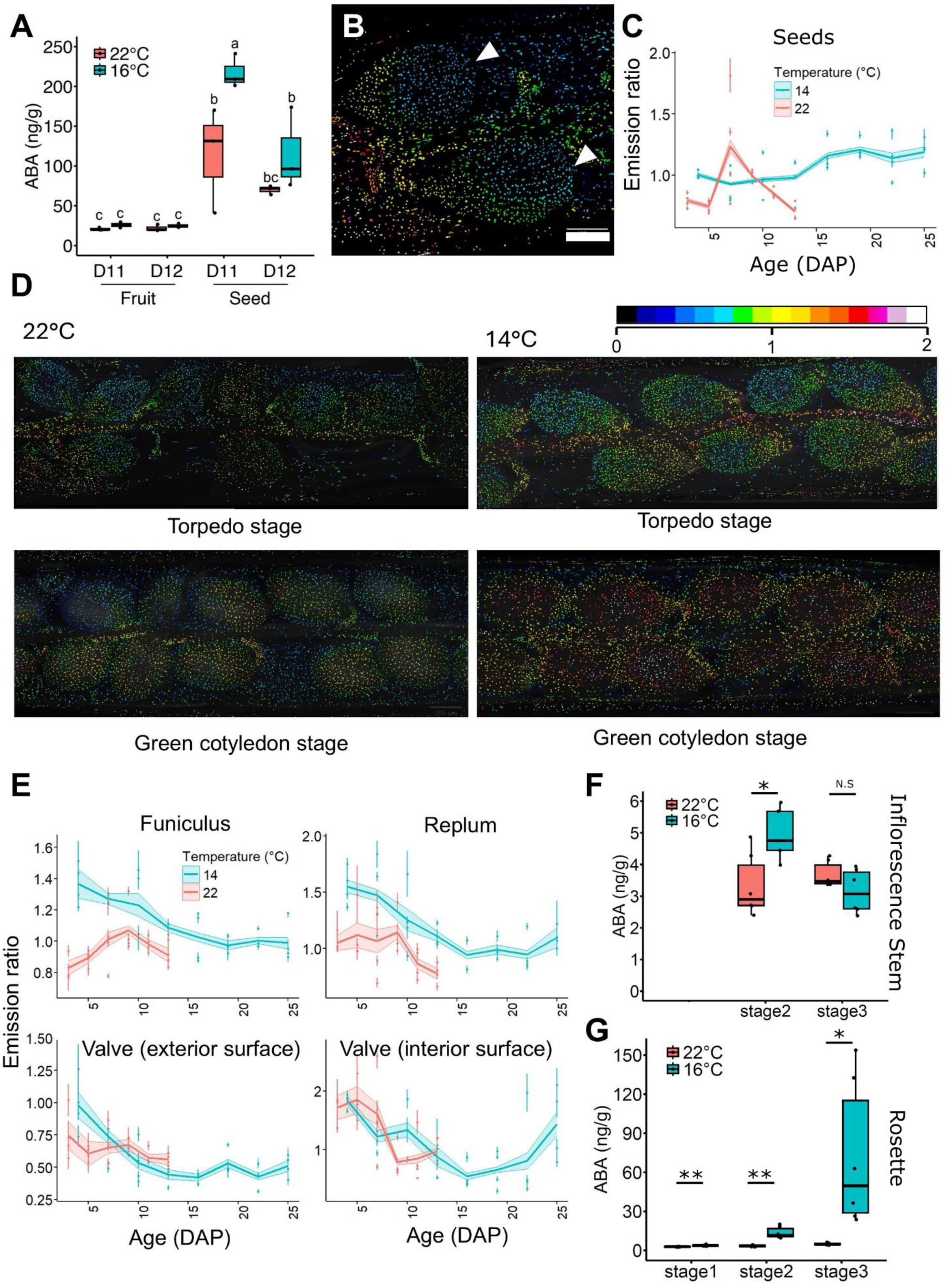
ABA levels, measured by LC-MS or biosensor imaging, show higher ABA in fruit and seed tissues at low temperatures and suggest a flux of ABA from fruit to seed tissues. **A.** LC-MS measurements of ABA in fruit (seeds removed) and seeds at 22°C or transferred to 16°C at 10 days after pollination. (D = Days after Pollination). Significant differences are shown by Tukey post hoc test (n = 3, P < 0.05). **B.** Example biosensor image of two seeds at 14°C 3 DAP attached to the replum showing a gradient of ABA from fruit to seed. White arrows indicate the distal pole of the seeds away from attachment to placental tissues. Scale bar 100µM. nuclei are false coloured by emission ratio as indicated from 0 to 2 with higher emission ratios corresponding to higher relative ABA levels. **C.** Quantified ABACUS2 emission ratios in seed tissues in a developmental series at low and high temperature with given ages in Days After Pollination (DAP). Start and end points at both temperatures are equivalent developmental stages. For C and E, individual points and error bars are the mean and 95% confidence interval of all nuclei within one biological replicate. Trendline and band around trendline indicate the mean and 95% confidence interval of the four biological replicates indicated for each combination of stage, tissue type, and temperature **D.** Example ABACUS2 biosensor images showing sections of fruits and seeds at low and high temperature at bent cotyledon and torpedo stage as indicated. False coloured nuclei by emission ratio are overlaid on brightfield images in grey. **E.** Quantified ABACUS2 emission ratios in different fruit tissues as indicated in a developmental series at low and high temperatures. **F, G.** LC-MS measurements of ABA levels in inflorescence stem (F) and rosette leaves (G) at different developmental stages at low and high temperatures (Figure S4). Significant differences are shown by t.test (n = 6, P < 0.05).

In seeds set in glasshouse conditions, a characteristic pattern of ABA accumulation occurs, peaking around the onset of seed maturation (Karssen et al., 1983, Kanno et al., 2010). We could observe similar dynamics with ABACUS2 at 22°C (Figure 6C, D) demonstrating that ABACUS2 faithfully reports ABA accumulation in seed tissues. In contrast, at 14°C ABA levels in seeds were already elevated early in seed development and continued to increase until maturity. In the fruit tissue, the effect of temperature depended on tissue type (Figure 6D, E): low temperature increased ABA levels primarily in the funiculus and in the replum in addition to the seeds themselves, showing that low temperature promotes higher ABA levels in the placental and vascular tissues connecting seeds to the mother plant. Tissues close to the abaxial epidermis of the valve showed high but temperature-independent ABA levels, whereas the adaxial epidermis had low levels of ABA in general. So in fruit tissues, low temperature effects on ABA levels are tissue-specific and consistent with ABA delivery through the vascular system from the mother plant into seeds.

In *lhp1-6* mutants, ABA biosynthetic gene expression is strongly down regulated in fruits but the same is not true in low temperature fruit tissues (Figure 2H). In fruit cultures temperature has a weaker effect on seed dormancy than that observed in inflorescence culture or whole plants (Figure S5) suggesting that ABA might arrive in fruits from more distant tissues of the mother plant. Consistent with this hypothesis we found that during flowering and seed set low temperatures increased ABA levels in the rosette leaves, but not so clearly in inflorescences where steady state ABA levels were very low (Figure 6F, G). Taken together our data reveal that temperature changes experienced by the mother plant are sensed via a nitrate response pathway and that maternal environmental information is communicated trans-generationally via the hormone flux of ABA from the mother plant into seeds.

## Discussion

Our data show that during reproductive development, plants form a memory of their climate history which is reflected in the germination properties of seeds. It has long been speculated that the effect of climate on seed dormancy is inherited from the mother plant (Roach and Wulff, 1987) but proof has been elusive and the mechanism has remained unclear. We show here that progeny dormancy responds to temperature changes in the mother plant but not the father (Figure 1B) and that progeny dormancy is prevented in mutants that affect ABA accumulation in seeds only when they are homozygous in the mother plant (Figure 1C, Figure 5D). We show that seeds must be attached to the mother plant for low temperatures to induce dormancy (Figure S2). Although many hormones affect seed dormancy and germination, we show that ABA applied to developing fruits accumulates in seeds and is sufficient to induce seed dormancy. We also show that maternal ABA levels during reproductive development respond strongly to small changes in ambient temperature and that this ABA is necessary and sufficient to induce progeny dormancy (Figure 5A). This cannot be easily detected in whole fruits because of varying ABA responses to temperature in different tissues, demonstrating the value of cell- and tissue-specific hormone analysis using biosensor technologies (Rowe et al., 2023). Long distance ABA transport plays other roles in plant response to the environment including root and shoot development and seed filling responses to environmental stresses (Qin et al., 2021; Zhang et al., 2023) suggesting a role in coordinating development and metabolism across distant tissues.

*Lhp1* mutants show a constitutively active primary nitrate response showing that *LHP1* is required for repression of the primary nitrate response in Arabidopsis: more surprising is that *lhp1* mutants accumulate nitrate (Figure 2). Our data show that temperature and soil nitrate activate a common response pathway during reproductive development and that at the physiological level nitrate can substitute for warm temperatures and vice versa in the control of progeny dormancy (Figure 2, 3). Furthermore, even moderate warming by a few degrees Celsius is sufficient to promote the PNR in plants and a change in tissue nitrate levels. Interestingly, a recent study also shows that nitrate affects high temperature stress responses in roots (Lee et al., 2024) suggesting that controlling nutrient use is a universal feature of plant responses to temperature changes. Climate adaptation is known to be mediated by the activity of natural selection on genetic and epigenetic variation in plants (Dubin et al., 2015). However, we reveal that short term changes in climate experienced by one generation can have immediate effects on phenotypes of the next generation via hormone transport. Thus, trans-generational hormone signalling is an alternative overlooked transmission mechanism and may play a role in re-establishing parental epigenetic states in progeny.

## Acknowledgements

The authors would like to thank James Rowe and Alexander Jones (Sainsbury Laboratory, Cambridge University) for the ABACUS2-400n seeds and invaluable advice for optimising our biosensor imaging and image processing, Eva Wegel and Sergio Lopez (JIC) for help with light microscopy, and John Innes Centre Horticultural Services and Lab Support. LHP1-GFP was given to us by Caroline Dean’s group. This work was supported by Biotechnology and Biological Sciences Research Council Grants BB/T003030/1 and BB/X015793/1 to S.P. and a John Innes Foundation studentship to W.B. The snRNA-seq experiment was delivered via Transformative Genomics the BBSRC funded National Bioscience Research Infrastructure (BBS/E/23NB0006) at Earlham Institute by members of the Single-Cell and Spatial Analysis and Core Bioinformatics Groups (V.K., A.G., A.L. and I.M.).

## Methods

### Resource availability

Further information and requests for resources and reagents should be directed to and will be fulfilled by the Lead Contact, Steven Penfield. The plasmids, transgenic plants and unique reagents generated in this study are available from the Lead Contact with a completed Materials Transfer Agreement.

### Data and code availability

All the bulk RNA-seq and single nuclei RNA-seq data generated from this study have been deposited in the Gene Expression Omnibus under accession code GSE278759 and GSEXXX.

### Plant materials and growth conditions

All Arabidopsis *thaliana* mutants used in this study have been previously described, including *lhp1-6* (SALK_011762) (Berry et al., 2017) and *lhp1-3* (Larsson et al., 1998), *aba2-1* (CS156) (González-Guzmán et al., 2002), *cyp707a2-1* (SALK_072410) (Kushiro et al., 2004). All mutants are in Col-0 background. pLHP1-LHP1-GFP/*lhp1-6* line was generated by crossing LHP1-eGFP/FRI^sf2^ (Berry et al., 2017) with Col0. pEPR1-LHP1-GFP/*lhp1-6* lines were generated by floral dipping with *Agrobacterium tumefaciens.* EPR1 promoter (2086bp) and LHP1 coding sequencing region (1356bp) was amplified by PCR, then assembled using Golden Gate with eGFP. The constructs were transformed into *lhp1-6* plants.

Arabidopsis plants were directly grown in long days (16h light at 22°C, 8h darkness at 20°C) till bolting. Then plants were either maintained in 22°C day/20°C night or transferred to 16°C day/14°C or 14°C day/14°C night. All mutants and wildtype were grown side-by-side for each experiment.

### Seed dormancy assay

For seed production, plants were grown in 16 hour light/8 hour dark for seed dormancy assays at either 22°C, 16°C or 14°C as indicated. Seed dormancy assays were performed by sowing freshly harvested seed onto 0.9% water agar plates. Seed batches from individual mother plants were used as biological replicates, at least 6 biological replicates of ∼100 seeds were used. Seed germination was scored by radicle emergence at 7 days after sowing. All statistical analyses in this study were performed using R (http://www.R-project.org/). For multiple tests, two-way ANOVA with post hoc test was applied using a threshold of p-value<0.05. All the other tests were performed using two-tailed Student’s t-test. The %germination data was arcsine transformed prior to statistical analysis.

### ABA measurement

ABA were measured as previously described (Chen et al., 2023). Different tissues were ground and extracted overnight at 4°C with 99 : 1, isopropanol : acetic acid. d6-ABA (OlChemIm) were added as internal standards. Supernatant was collected after centrifugation before drying in an Evaporator. The dried extracts were resuspended in methanol and filtered through 0.22-μm Corning® Costar® Spin-X® Plastic Centrifuge Tube Filters (Sigma). The solution was injected and analysed on an ultraperformance liquid chromatography-mass spectrometry (UPLC-MS) system. For the d6-ABA transport experiment, internal standards were not added and standard curves with known concentrations of both unlabelled ABA and d6-ABA were used.

### Nitrate measurement

Plant material was flash frozen and ground to a powder using a QIAGEN TissueLyser with 3mm tungsten carbide beads (QIAGEN). 10-100mg of powder were weighed out and nitrate extracted in ddH_2_0 for 1 hour at room temperature on an orbital shaker. Supernatant was collected after centrifugation and filtered through 40μM Corning Cell Strainers. The filtered supernatant was analysed for nitrate+nitrite and nitrite using an AQ300 Discrete Analyser (SEAL Analytical) by the University of East Anglia Elemental Analysis Platform.

### In vivo ABA detection using the ABACUS2 Biosensor

The ABACUS2 400n Biosensor (Rowe *et al*., 2023) was used for *in vivo* relative ABA measurements. Imaging was performed using a Zeiss AxioZoom.v16 with 1x objective lens. Excitation used a Lumencor SPECTRA3 Light Engine with blue LED at 423-452nm and 438/29 filter at 30% intensity to excite CFP. Emission was collected for CFP with filter 525/50 using the Zeiss 47 HE CFP filter cube representing the unbound biosensor and for YFP another Zeiss 47 CFP filter cube modified with the 544/24nm bandpass filter for YFP emission. Prior to imaging, the first flowers were screened for high expression of nuclear localised YFP signal and only those with signal readily detected by epifluorescence were used. All images were taken of live material starting within 5 minutes of removal of fruits from plants. The upper part of both valves was removed by dissection with a fine needle and the sample mounted in perfluorodecalin for live imaging.

Analysis was performed in the FIJI distribution of ImageJ (Schindelin *et al*., 2012) using the FRETENATOR2 Plugin (Rowe *et al*., 2023). Segmentation was performed using the FRETENATOR2 Segment and Ratio function using the CFP excitation, CFP emission channel for segmentation and excluding ROI below 50 or above 1000 pixels in size. For quantification by tissue, a representative tile for each image for four biological replicates per temperature and timepoint combination was processed using the ROI Labeller function.

### Fruit culture

Complete fruit culture medium is modified from that used by Creff *et al.,* (2023) for *in vitro* fruit culture. Complete fruit culture medium is 1/2 strength Murashige and Skoog media including vitamins and Gambourg’s B5 basal salt mixture, with 1% sucrose, and 1/1000 Plant Preservative Mixture (Apollo Scientific) in ddH_2_O and autoclaved. For experiments with modified nitrate level, Ammonium Nitrate and Potassium Nitrate is adjusted in proportion from the complete fruit culture medium. ABA is added from a 10mg/ml stock in methanol immediately before use to indicated concentrations. For fruit culture, flowers are marked on the day of fertilisation and fruits at three days after pollination are removed and placed in 1.85ml of fruit culture medium in 2ml centrifuge tubes. The pedicels are then re-cut once submerged using a fine needle to prevent air bubble formation blocking vasculature and placed back in growth cabinets in transparent tube racks. For inflorescence culture, including the d6-ABA transport experiment, 40ml of culture medium is put in 50ml centrifuge tubes and sealed with parafilm. Young inflorescences where the oldest buds have been fertilised within 24 hours are cut from the plant and pushed through a hole in the parafilm so that the lower part is submerged but all fruits/flowers are above the medium containing d6-ABA.

### RNA-seq and data analysis

Arabidopsis plants were grown in 22°C until bolting, then transferred to 16°C or 22°C as indicated. The Day 0 (0DAP) was identified by the first emergence of petals from the buds. At bent-cotyledon stage, Col16 (14-15DAP), Col22(6-7DAP), *lhp1*(14-15DAP) siliques were manually dissected under microscope. Separated fruits and seeds from 3-5 siliques were collected as one biological replicate. RNA was extracted using RNeasy Plant Mini Kit (Qiagen, 74904). Fruits and seeds RNA samples were sequenced at BGI (China) using DNBSEQ, 100-bp paired-end sequences with minimum 25 million reads acquired per sample.

Raw RNA-seq reads were trimmed using cutadapt-1.9.1 (Martin, 2011) and mapped to *Arabidopsis thaliana* TAIR10 reference genome using STAR-2.5.a (Dobin et al., 2013). featureCounts (Liao et al., 2014) was used to count the numbers of reads mapped to each gene. Sense reads were selected for downstream analysis. Differentially expressed genes (DEGs) was calculated using edgeR (Robinson et al., 2010) with the threshold of FDR<0.05 and fold change≥2.

### GO analysis

The GO analysis was performed using ‘topGO’ (Alexa & Rahnenfuhrer, 2024). Briefly, differentially expressed gene lists of interest were analysed using the topGO runTest function, with algorithm = ‘elim’ and statistic =‘fisher’. GO annotations were obtained from plants_mart at plants.ensembl.org using the ‘biomaRt’ package (Durinck et al., 2005). All genes with detectable expression in bulk RNA-seq were set as the background set of genes.

### ChIP-seq data analysis

The LHP-GFP ChIP-seq data (Veluchamy et al., 2016) was re-analyzed. Raw reads were trimmed using cutadapt-1.9.1 (Martin, 2011) and mapped to *Arabidopsis thaliana* TAIR10 reference genome using bowtie2 (Langmead and Salzberg, 2012). Only uniquely mapped reads were kept for downstream analysis using Samtools-1.9 and Sambamba-6.7 (Li *et al*., 2009; Tarasov *et al*., 2015). bigwig files were calculated using deepTools-3.1.1 (Ramirez *et al*., 2014) with a bin size of 50bp, before visualization in IGV -2.12.3 (Robinson *et al*., 2011). Peaks were called using MACS3 (Zhang *et al*., 2008), based on the threshold of FDR<0.05.

### Single nuclei preparation, library construction and sequencing

Col-0 and *lhp1* plants were grown at 22°C until bolting, then transferred to 16°C. The Day 0 (0DAP) was identified by the first emergence of petals from the buds. *lhp1* plants were grown in 16°C for 14 days (*lhp1* 16°C sample). Col-0 plants were grown at 16°C for 13 days, with half plants moved to 22°C for 1 day (Col-0 22°C sample), while another half plants left at 16°C for 1 day (Col-0 16°C sample).

20-30 siliques from each sample were placed on a petri dish, and chopped in 3min in 2ml ice-cold nuclei extraction buffer (Marand et al., 2021), as one biological replicate. Two biological replicates were prepared for each sample. The nuclei solution was further filtered through 30 um cell strainers (Sysmex). After staining with 10μl of 1mg/ml DAPI, 150K nuclei were sorted into 20μl collection buffer (Marand et al., 2021) on a BD FACSAriaIII flow cytometer. Nuclei were sorted with a 70 micron nozzle and 4-Way Purity precision setting with 405nm laser excitation, 450/50nm emission detection filter and gating strategy as depicted in representative plot (Figure S6B) sorting diploid and triploid nuclei. Sorted nuclei were counted and quality checked by microscopy (Figure S6A), and 16k nuclei loaded into Chromium X Controller (10X Genomics) by the Single-cell and Spatial Analysis platform at the Earlham Institute (Norwich, UK), with a target recovery of 10K. Libraries were further constructed with reagents from Chromium Single Cell v3.1 reagent kit (10X Genomics) according to the manufacturer’s instructions and sequenced at the Earlham Institute.

### Single nuclei RNA-seq data analysis

The raw snRNA-seq dataset was analysed using Cell Ranger-7.2.0 (10X Genomics) for early-stage samples and Cell Ranger-8.0.0 for late-stage samples. The Arabidopsis TAIR10 reference genome and Araport11 GTF annotation files were used for downstream analysis, according to the default cellranger mkref and count commands. The gene-cell matrices were loaded into the Seurat Package-5.1.0 (Hao et al., 2023), with min.cells = 3 and min.features = 200. Then the cells were further filtered with >150 nFeature_RNA, <4000 nFeature_RNA, <5% mitochondrial sequences and <20% chloroplast sequences. The six Col16, Col22 and lhp16 samples from early- or late-stage were integrated for downstream analysis. Col22 replicate2 sample was excluded due to it’s deviated distribution from other samples after integration (Fig.S7, Table S7). After comparing a series of resolution, we selected resolution =0.1 for further analysis (Figure S3A) To separate fruit and seed cell types, we first compared the bulk fruit and seed RNA-seq data using edgeR, to generate the fruit-specific genes (Table S3), with FDR<0.01 and Fold change>10. Then we matched these fruit-specific genes to single nuclei RNA-seq data to identify fruit and seed cell clusters. All tissue/cell-type markers were obtained from literatures, including funiculus, phloem, xylem, seed coat, endosperm, embryo, epidermis and cortex (Table S3).

### Statistical Analysis

All statistical analyses in this study were performed using R (http://www.R-project.org/). For multiple tests, one-way ANOVA with post hoc test was applied using a threshold of p-value<0.05. All the other tests were performed using two-tailed Student’s t-test. The %germination data was arcsine transformed prior to statistical analysis.

## Supplementary Figures S1-S6

**Figure S1.**
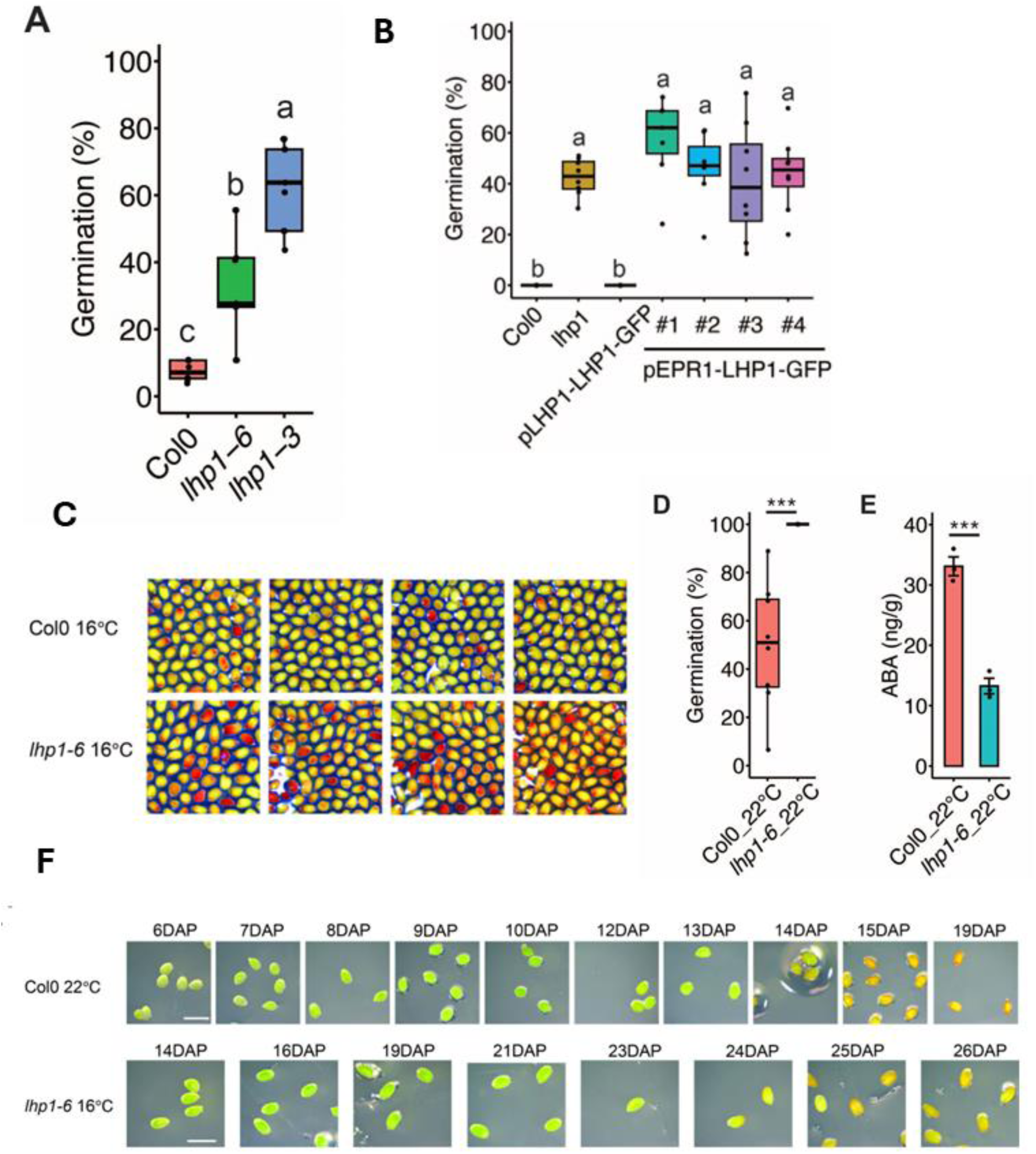
**A.** Comparison of *lhp1-3* and *lhp1-6* alleles to Col-0. **B.** Complementation of *lhp1-6* with pLHP1:LHP1-GFP restores the WT seed dormancy phenotype. Complementation with zygotic expression using pEPR:LHP1-GFP does not restore the WT phenotype. **C.** Permeability of *lhp1-6* and WT Col-0 seeds. **D.** Germination of *lhp1-6* and WT Col-0 set at 22°C and **E.** corresponding ABA levels. **F.** Images of seed morphology during development, corresponding to germination data in Figure 1A.

**Figure S2.**
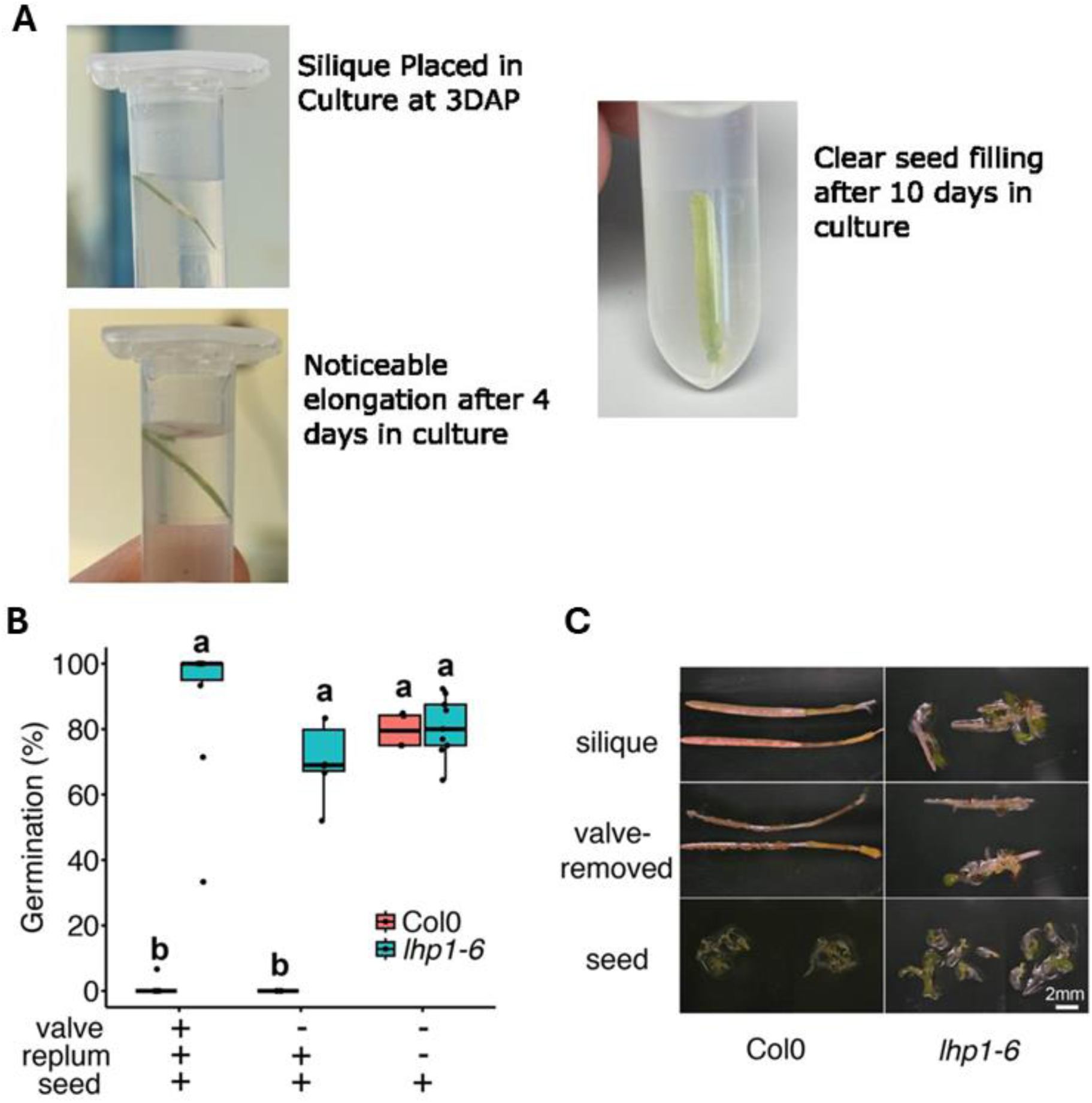
**A.** Images showing the process of viable seed set in fruit culture. **B.** and **C.** LHP1 prevents pre-harvest sprouting. Attachment to the replum with or without valves is sufficient to prevent germination in fruit culture in WT Col-0. Premature germination occurs in *lhp1-6*.

**Figure S3.**
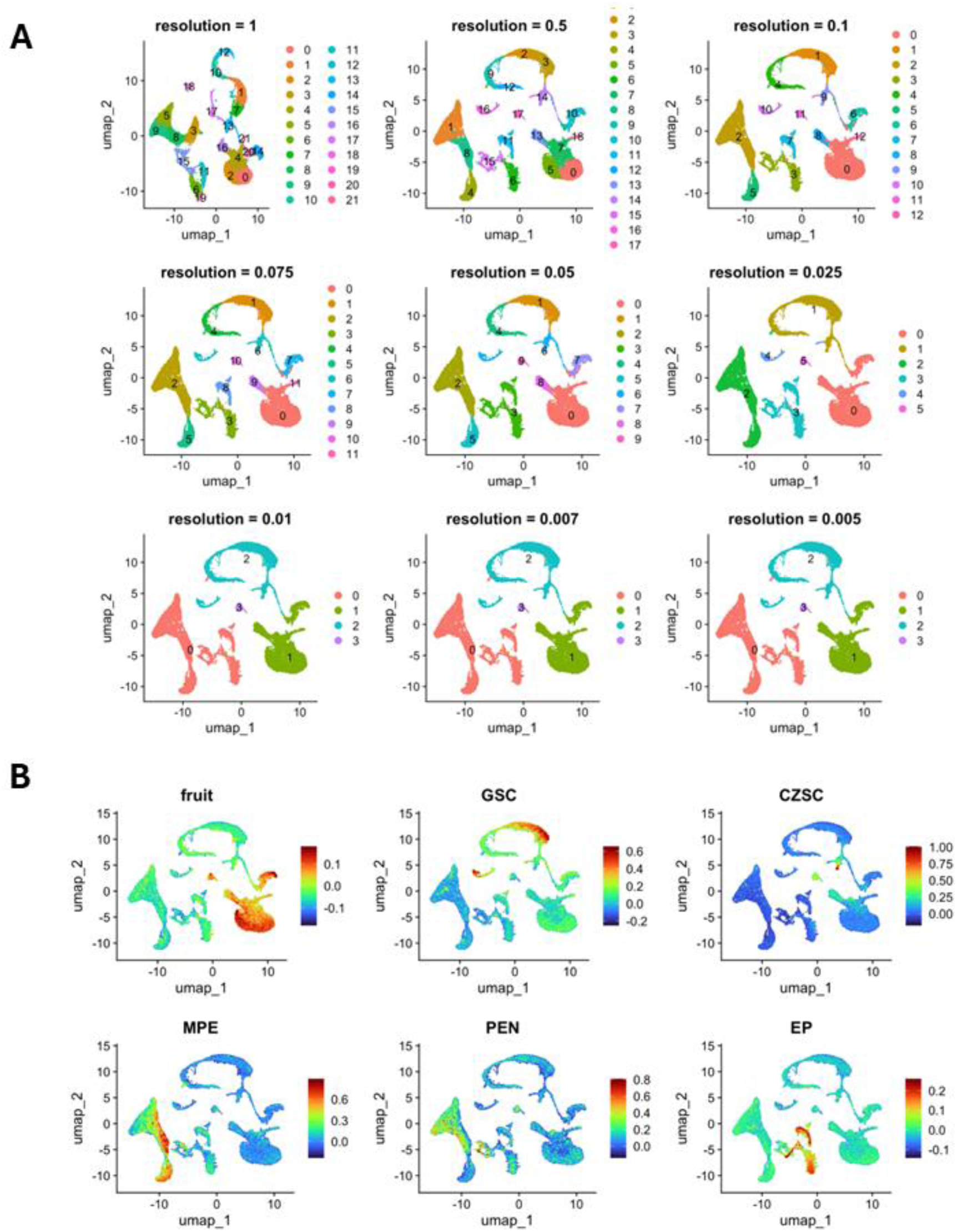
**A.** Clustering at varied resolution indicates broader tissue types as well as more specific cell types. Resolution indicates Seurat clustering parameter. **B**. Genes enriched in expression in fruits or in different regions of seeds plotted on UMAP plots as in Figure 3A provides additional evidence to support our annotated tissue types. (GSC = General Seed Coat, CZSC = Chalazal Zone Seed Coat, MPE = MicroPylar Endosperm, PEN = Peripheral Endosperm, EP = Embryo Proper).

**Figure S4.**
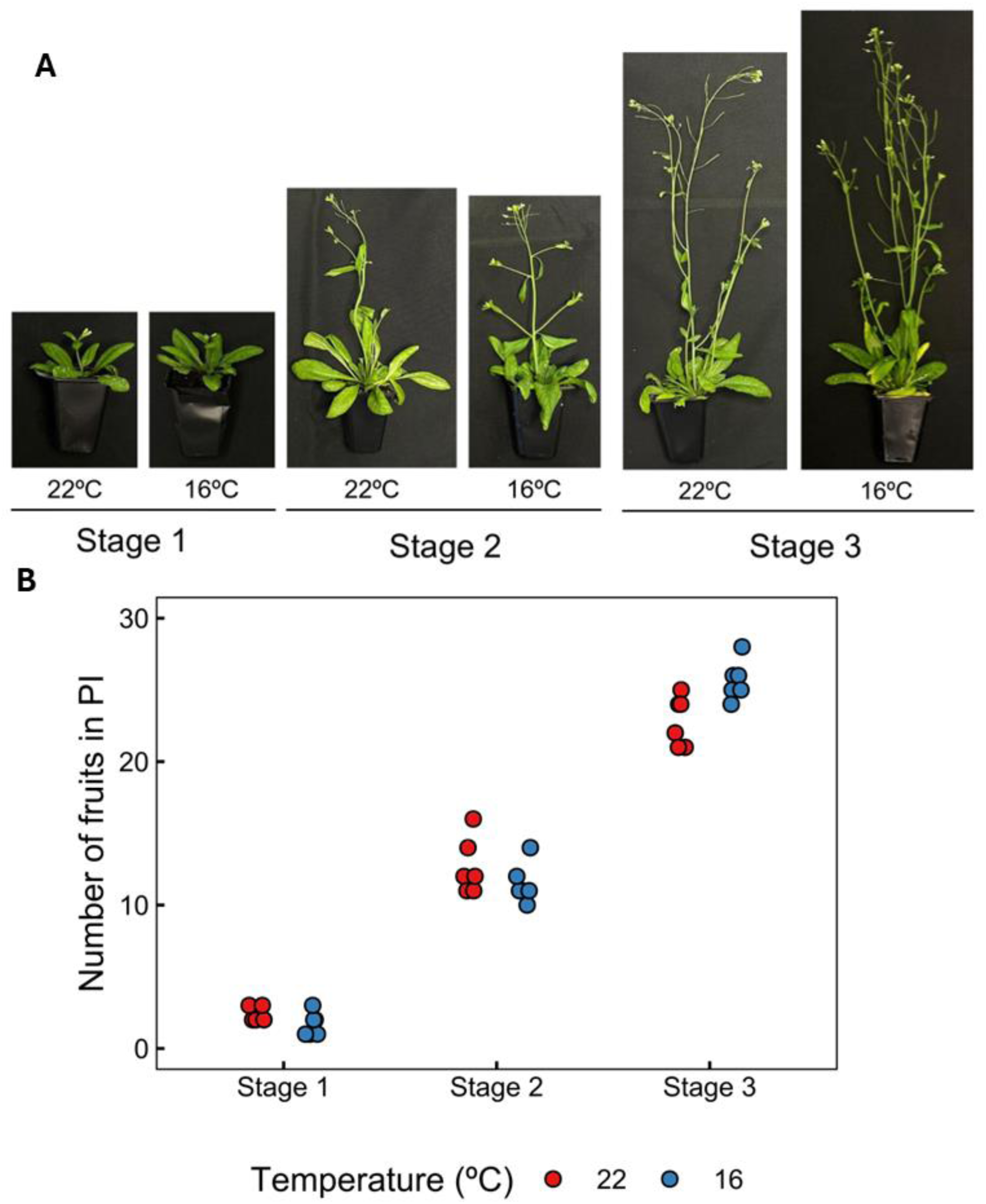
Developmental stages shown in Figure 4F&G and Figure 5 F&G. **A.** Examples of plants grown at low and high temperature corresponding to stages. **B.** Numbers of fruits on Primary Inflorescence (PI) used to match developmental stages.

**Figure S5.**
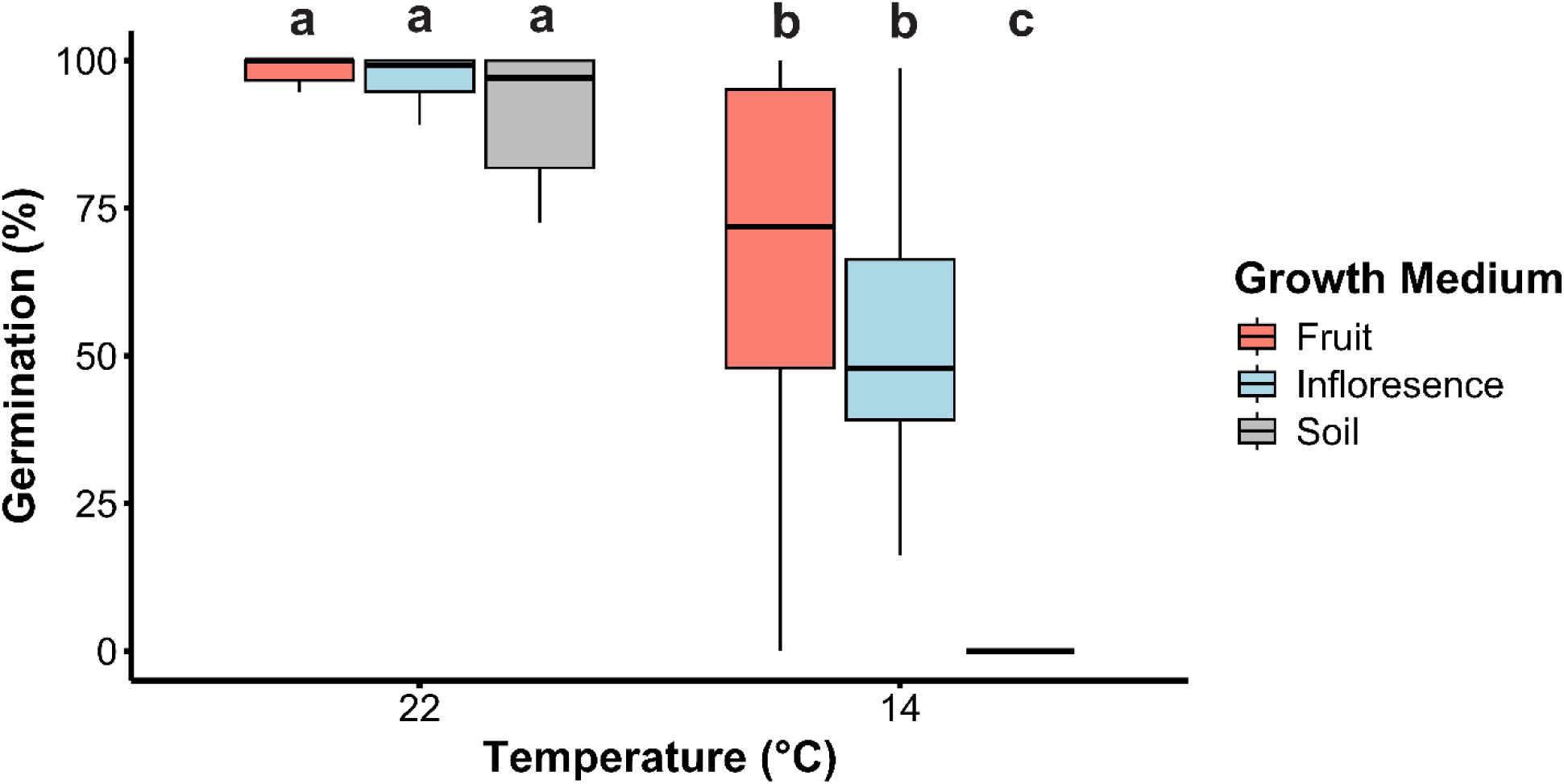
Germination of WT Col-0 seeds produced from fruit culture, inflorescence culture, or on soil-grown plants at the indicated temperatures.

**Figure S6.**
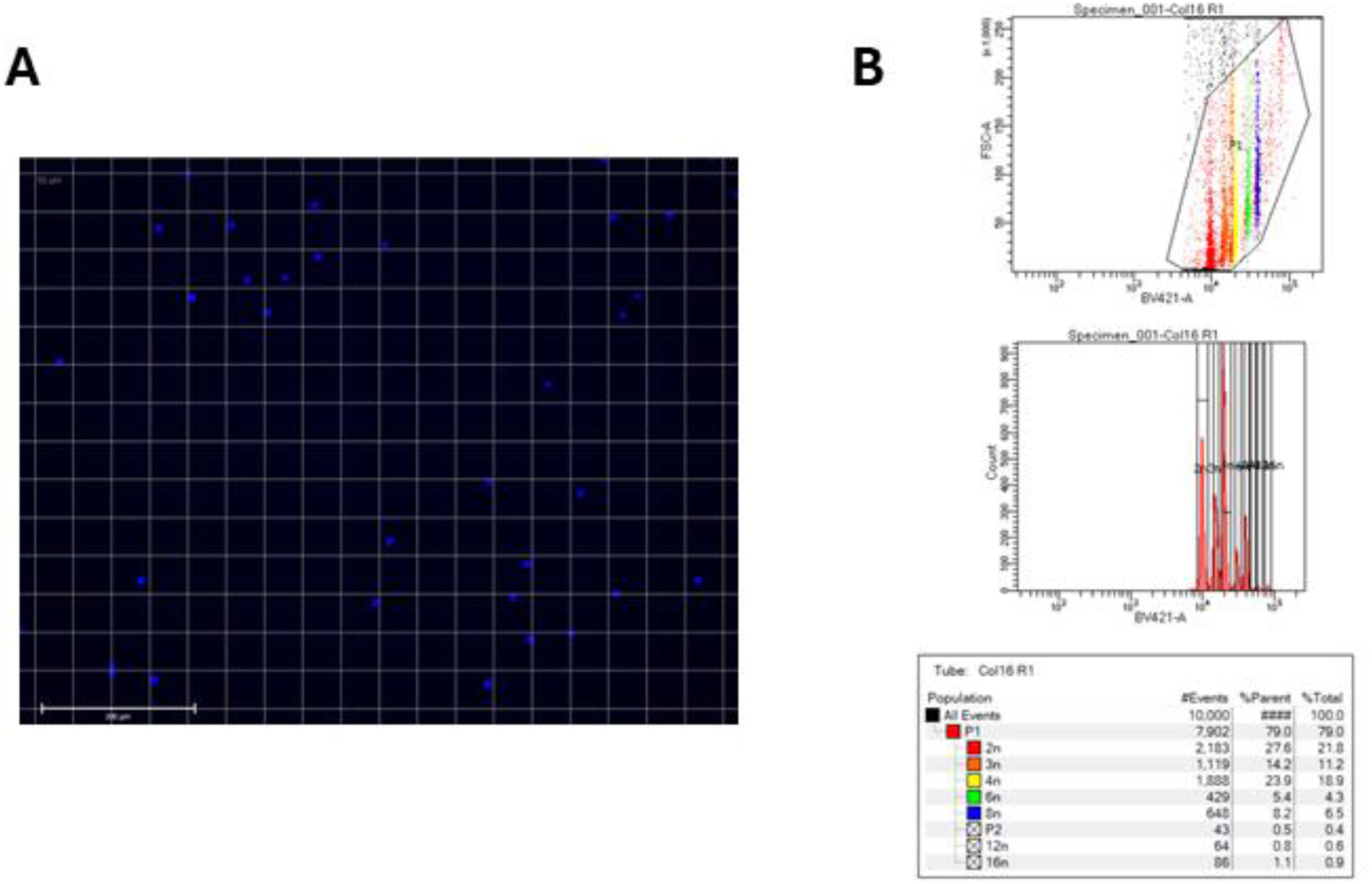
**A.** Image of DAPI stained nuclei after FACS sorting showing healthy intact nuclei. **B.** Example of a sort plot showing parameters used to isolate nuclei for snRNA-sequencing. DAPI 405nm laser excitation with 450/50nm band pass filter detection.

**Figure S7.**
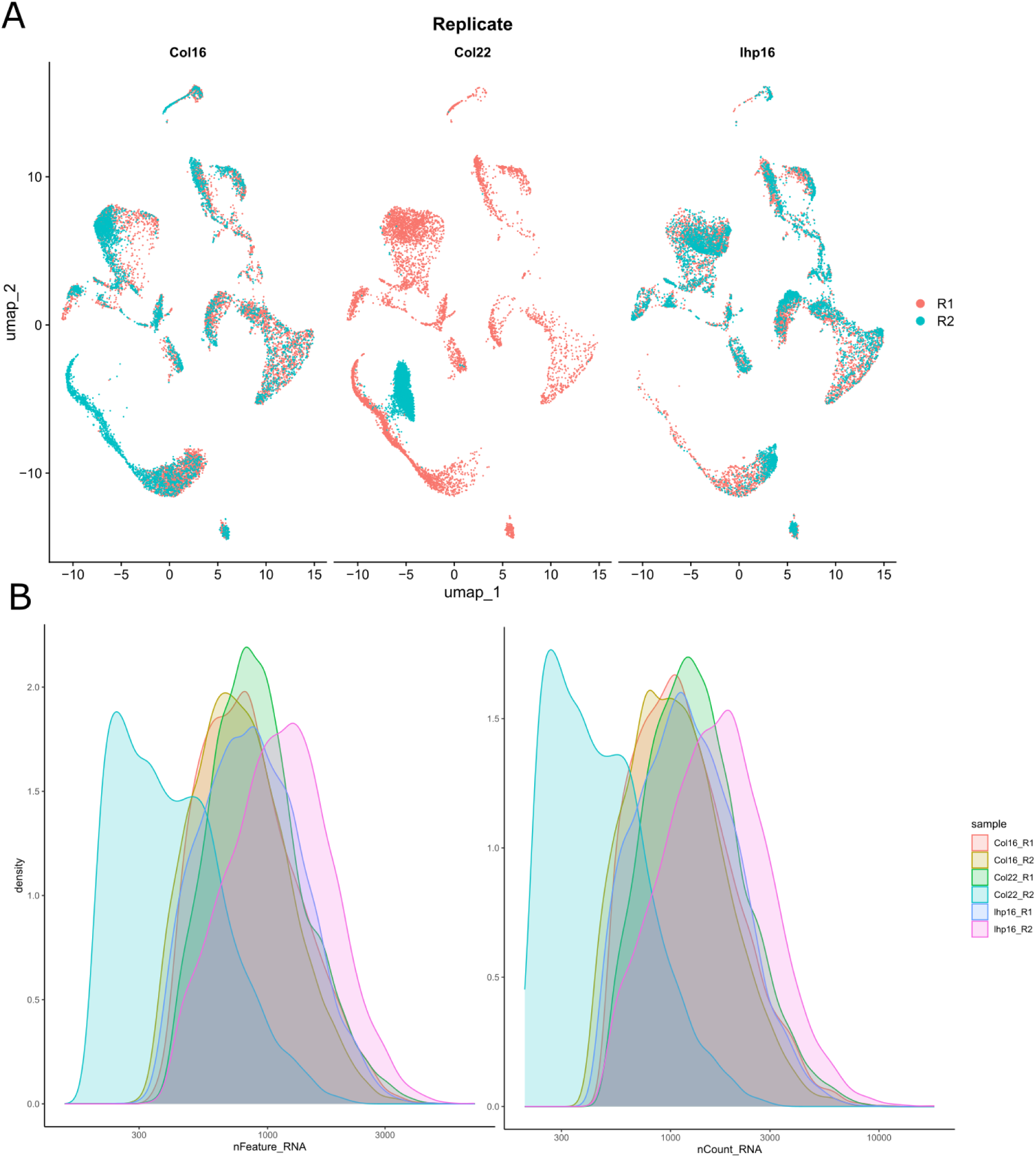
Col-0 22°C Replicate 2 (Col22_R2) was excluded from downstream analysis. **A.** All detected nuclei in Col22 R2 cluster together and away from other nuclei. **B.** Nuclei from Col22 R2 have noticeably fewer unique genes (nFeature_RNA) and few RNA molecules (nCount_RNA) compared to all other samples.

